# Terminal differentiation of villus-tip enterocytes is governed by distinct members of Tgfβ superfamily

**DOI:** 10.1101/2022.11.11.516138

**Authors:** Linda Berková, Hassan Fazilaty, Qiutan Yang, Jan Kubovčiak, Monika Stastna, Dusan Hrckulak, Martina Vojtechova, Michael David Brügger, George Hausmann, Prisca Liberali, Vladimir Korinek, Konrad Basler, Tomas Valenta

## Abstract

The protective and absorptive functions of the intestinal epithelium rely on differentiated enterocytes in the villi. The differentiation of enterocytes is orchestrated by sub-epithelial mesenchymal cells producing distinct ligands along the villus axis, in particular Bmps and Tgfβ. Here we show that individual Bmp ligands and Tgfβ drive distinct enterocytic programs specific to villus zonation. Bmp4 is expressed mainly from the center to the upper part of the villus, and it activates preferentially genes connected to lipid uptake and metabolism. In contrast, Bmp2 is produced by villus-tip mesenchymal cells, and it influences the adhesive properties of villus-tip epithelial cells and the expression of immunomodulators. Hence, Bmp2 promotes the terminal enterocytic differentiation at the villus-tip. Additionally, Tgfβ induces epithelial gene expression programs similar to that triggered by Bmp2. The inhibition of Bmp receptor type I *in vivo* and using intestinal organoids lacking Smad4 revealed that Bmp2-driven villus-tip program is activated by a canonical Smad-dependent mechanism. Finally, we established an organoid cultivation system that enriches for villus-tip enterocytes and thereby better mimics the cellular composition of the intestinal epithelium. Altogether our data suggest that not only Bmp gradient, but also the activity of individual Bmp drives specific enterocytic programs.

## Introduction

The small intestinal epithelium consists of diverse types of differentiated cells that are important for a wide range of functions including absorption of nutrients and protection of the body from the intestinal content. The differentiated cells arise in the crypt from the intestinal epithelial stem cells (IESCs), respectively from their progeny, transit amplifying cells^1,2^. The differentiated cells exit the crypt, migrate upwards along the villus and are eventually shed into the intestinal lumen from the villus tip. It is thought that various stromal signals along the crypt-villus axis orchestrate the differentiation process. The details of this complex process have only just begun to be elucidated. Since dysregulation underpins many intestinal diseases, it is imperative to define the output of the various signals provided along the crypt-villus axis and their role during the intestinal homeostasis.

Enterocytes are the most abundant differentiated intestinal epithelial cells with a life-time of approximately 4 days^3,4^. Besides absorbing nutrients, enterocytes constitute a physical and biochemical barrier, produce antimicrobial peptides and can act as immunomodulators^4,5^. Enterocytes display a zonated gene expression pattern along the crypt-villus axis^5^. Enterocytes at the bottom of the villus express antimicrobial programs, while in the center and top of the villus they sequentially activate genes connected to the absorption of amino acids, carbohydrates and lipids. In addition, as they migrate towards the villus-tip, enterocytes upregulate genes related to cell adhesion and immunomodulation^4,5^.

The zonated gene expression signatures, as well as the fate of enterocytes, are most likely governed by subepithelial mesenchymal cells producing morphogen gradients. In the crypt, Pdgfra^low^ mesenchymal cells secrete Wnt ligands that support the renewal of IESCs and intestinal homeostasis^6–9^. Besides Wnts, they also produce inhibitors of Bone morphogenic proteins (Bmp) such as Gremlin possibly creating an insulating zone around the stem cells and proliferating progenitors, separating IESCs from differentiated cells. In the villus, Pdgfra^high^ mesenchymal cells express various Bmp ligands (further referred to as Bmps)^7,8^. Bmps belong to Transforming growth factor β (Tgfβ) superfamily of secreted morphogens. They can form homo- or heterodimers that signal via canonical Smad-dependent pathway or induce various non-canonical signalling routes^10,11^. In the intestine, canonical Bmp pathway directly restricts the stemness of IESCs and thus counteracts Wnt signaling^12^. Additionally, Bmps trigger the differentiation of epithelial cells, in particular enterocytes, along the crypt-villus. Bmp2 and Bmp4 are the two most abundant Bmp ligands in the intestine. The combination of Bmp2 and Bmp4 activates genes responsible for lipid uptake^13^. Of note, Bmp2 and Bmp4 do not form heterodimers inter se, thus they can also act differently^10^. However, the individual roles of Bmp2 vs. Bmp4 remain unknown. The transcriptional switch from carbohydrate to lipid metabolism (initiated either by Bmp2 or by Bmp4) occurs near the center of the villus and the resulting subtype of enterocytes are found also in the upper part of the villus, but not at the villus tip^13^. In the enterocytes, Bmps activate c-Maf, a master regulator of lipid and amino acid uptake^14,15^. Hence, Bmps trigger the differentiation of epithelial cells, in particular enterocytes, manifested as sequentially activated and zonated-gene expression along crypt-villus axis.

Although it is well-described that along the villus the features of the enterocytes gradually change, we still do not understand how exactly this process is regulated. Is it simply the vaguely described Bmp gradient that is responsible for this process? Why are then various Bmps expressed in the villus, if just one Bmp can form the gradient and guide the entire process? Are the individual intestinal Bmps redundant or interchangeable? Do they act distinctly? How do they contribute to unique features of villus tip enterocytes? To address these questions, we set out to study the role of distinct members of the Tgfβ superfamily, particularly Bmps, in regulating the differentiation of intestinal cell types and the functional zonation of the villus using in vitro organoids models and in vivo mouse models. We find Bmp2 and Tgfβ induce distinct villus-tip programs, while Bmp4 regulated the villus center genes. Further, we establish a new organoid culture system, enriched in villus-tip enterocytes.

## Results

### Villus mesenchymal cells induce differentiation of intestinal epithelial cells via Bmp

Villus mesenchymal cells corresponding to Pdgfra^high^ cells in the small intestine are expected to control differentiation of epithelial cells^7,8,16^. To understand how they influence the fate of the epithelial cells and to explore the role of factors secreted by them we generated an immortalized small intestinal Pdgfra^high^ cell line.

Despite the long term *in vitro* cultivation and immortalisation procedure (4-6 weeks) the immortalized villus mesenchymal cells (ivMCs) retained key features typical for *in situ* to Pdgfra^high^ cells of the small intestine such as the expression of Bmps and the absence of canonical Wnts (Fig. EV 1A,B). Hence, ivMCs cell line can serve as a proxy for to Pdgfra^high^ cells of the small intestine. In co-cultivation experiments ivMCs induced differentiation of freshly isolated intestinal crypts into secretory lineage such as goblet cells (marked by *Muc2*) and enterocytes (*Krt20, Alpi, Sim* - positive); the cell morphology also changed (Fig. 1A,B). Importantly, ivMCs promoted the terminal differentiation of enterocytes determined by the expression of villus-tip genes (Fig.1B). The induction of the villus-tip program could be partially abrogated by treatment with the Bmp type I receptor inhibitor LDN-193189, pointing to the role of Bmps in terminal enterocytic differentiation (Fig. EV 1C,D).

**Figure 1.**
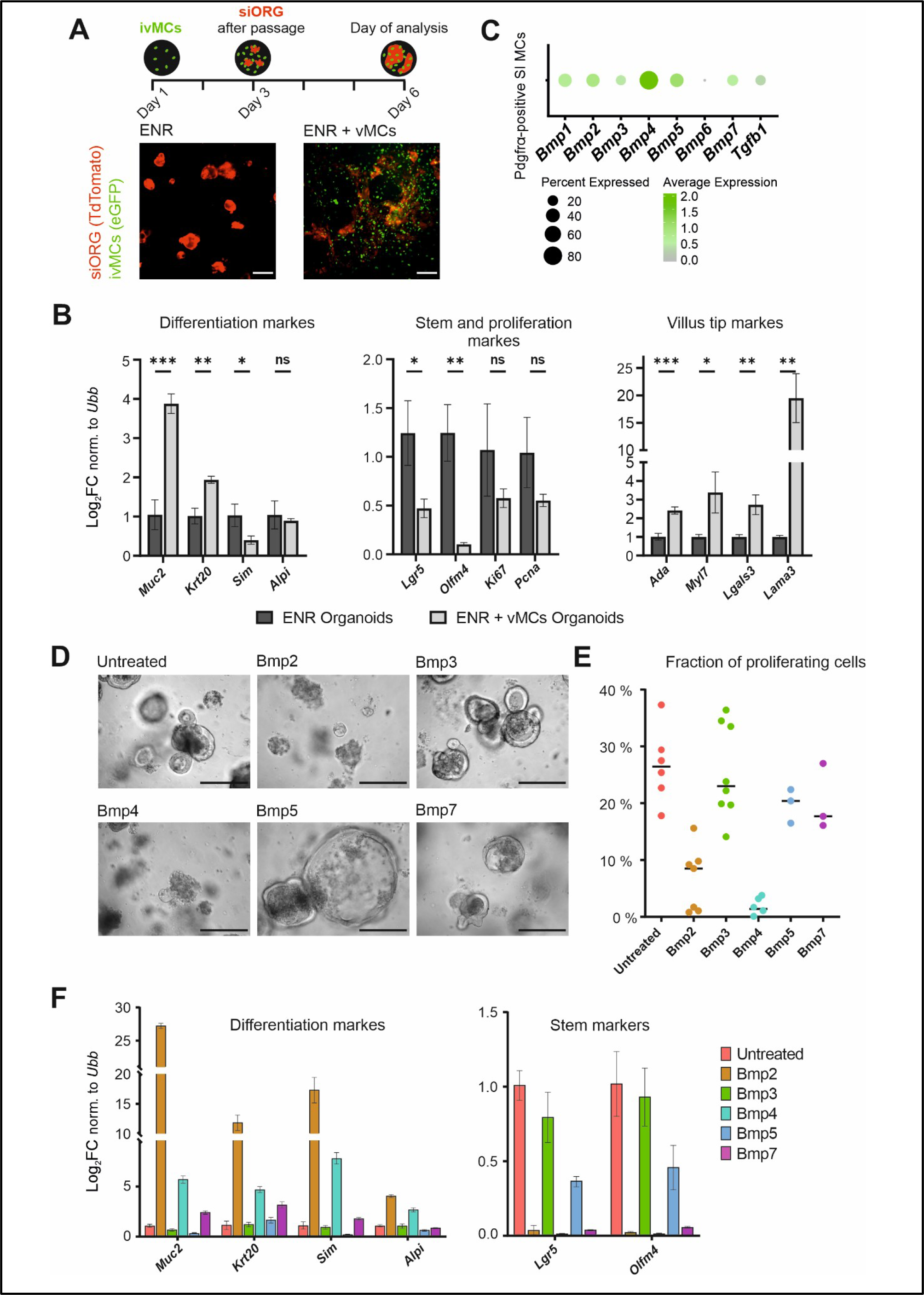
Bmp ligands secreted by villus Mesenchymal Cells are sufficient to boost intestinal epithelial differentiation. A Co-cultivation of intestinal organoids (siORG, marked by tdTomato) with immortalized villus Mesenchymal cells (ivMCs, expressing nuclear eGFP). ENR – organoids in cultivation medium (no ivMCs), ENR + ivMCs organoids in cultivation medium with ivMCs. (Native fluorescent microscopy. Scale bar, 200 μm). B ivMCs induce differentiation of co-cultured intestinal organoids as determined by qRT-PCR for differentiation markers (*Muc2*-goblet cells, *Krt20*-differentiated epithelial cells, *Sim* and *Alpi*-enterocytes), by reduced expression of stem cell (*Lgr5*, *Olfm4*) and proliferation genes (*Ki67*, *Pcna*) or upregulated markers of villus tip enterocytes. (Expression levels normalized to *Ubiquitin B* (*Ubb*), n=3, ENR parallel set as 1, error bars show sd, unpaired t-test, ns: p > 0.05, * p ≤ 0.05, ** p ≤ 0.01, *** p ≤ 0.001). C Expression of individual Tgfβ and Bmp ligands in Pdgfra mesenchymal cells of small intestine (SI). (Dot plot, size of the dot represents the percentage of the cells expressing the transcript, color indicates the expression level. Only ligands determined to be expressed are shown. Used dataset: GSM3747599) D The impact of individual Bmp on the growth of freshly isolated intestinal crypts. (Time point = after 96 hours in 500 ng/ml of indicated Bmp. Brightfield images. Scale bar, 200 μm). E Proliferation of freshy isolated intestinal crypts cultivated with indicated Bmps for 96 hours determined by quantification EdU incorporation (% of EdU positive cells). Each dot represents one cultured organoid. Average value for each treatment indicated by line. F Relative expression of differentiation and stem cells markers in intestinal crypts upon 96 hours cultivation with indicated Bmp (500 ng/ml). (qRT-PCR, expression levels normalized to *Ubiquitin B* (*Ubb*), n=3, error bars show sd, untreated parallel set as 1).

Re-analysis of published scRNAseq dataset revealed that sub-epithelial mesenchymal cells marked by Pdgfra in the small intestine display a complex expression pattern of distinct Tgfβ superfamily members (Fig. 1C). The striking complexity and diversity of Bmps secreted by sub-epithelial cells lead us to ask if individual Bmps have similar effect on epithelial cells, or if a particular Bmp has its specific role in the regulation of the intestinal differentiation.

To begin to address this question, we assessed the impact of individual Bmps on gene expression, morphology and cellular composition of organoids. Freshly isolated intestinal crypts were treated by selected recombinant Bmp ligands (Fig. EV 1E). After 96 hours (i.e. time corresponding to the average life of enterocytes in the villus) Bmp-treated organoids showed distinct changes in shape and survival (Fig. 1D), proliferation rate (Fig. 1E and Fig. EV 1F) and gene expression (Fig. 1F). The induced changes resembled those observed after ivMCs cocultivation, while Bmp2 and Bmp4 had the strongest effect. Based on expressed markers, Bmp2, Bmp4 and Bmp7 promoted differentiation to goblet cells (*Muc2*) and enterocytes (*Krt20, Sim, Alpi*). Additionally, the IESC pool was exhausted and proliferation was reduced, mirroring enhanced differentiation at the cost of IESCs (Fig. 1F).

### Individual Bmps activate distinct enterocytic differentiation programs

During their movement upwards, the enterocytes quickly changed their position and could be exposed to particular Bmp level or the particular Bmp ligands only for a short time^4^. To model short term Bmp exposure and to delineate Bmp-driven differentiation programs we treated intestinal crypts (72 after the isolation) with individual recombinant Bmps for 24 hours followed by RNA sequencing (Fig. EV 2A). Short-term treatment had less impact on the growth and viability of the organoids than 96 hours of treatment (Fig. EV 2B).

Principal component analysis of differentially expressed genes based on at least three independent experiments revealed individual Bmps had distinct effects (Fig. 2A). Bmp2 and Bmp4 treated samples grouped separately, suggesting specific targets for each of them. Bmp5 and Bmp7 treated samples grouped together between Bmp2 and untreated samples, pointing to similar but milder effect as compared to Bmp2. Bmp3 overlapped with untreated samples, indicating that Bmp3 had no effect on the organoids, most likely connected to its specific antagonistic role^10,17^ (Fig. 2A). The missing activation capability of Bmp3 was recently shown also for human intestinal organoids.^13^ Hierarchical clustering, based on the top 10 differentially expressed genes for each treatment, revealed that Bmp2 and Bmp4 treatment have different genetic outputs (Fig. 2B). This differential output is not related to differences in the specific activity of the recombinant Bmps since Bmp2 and Bmp4 activated the generic Bmp targets Id1 and Id3 to a similar extent (Fig. EV 2C).

**Figure 2.**
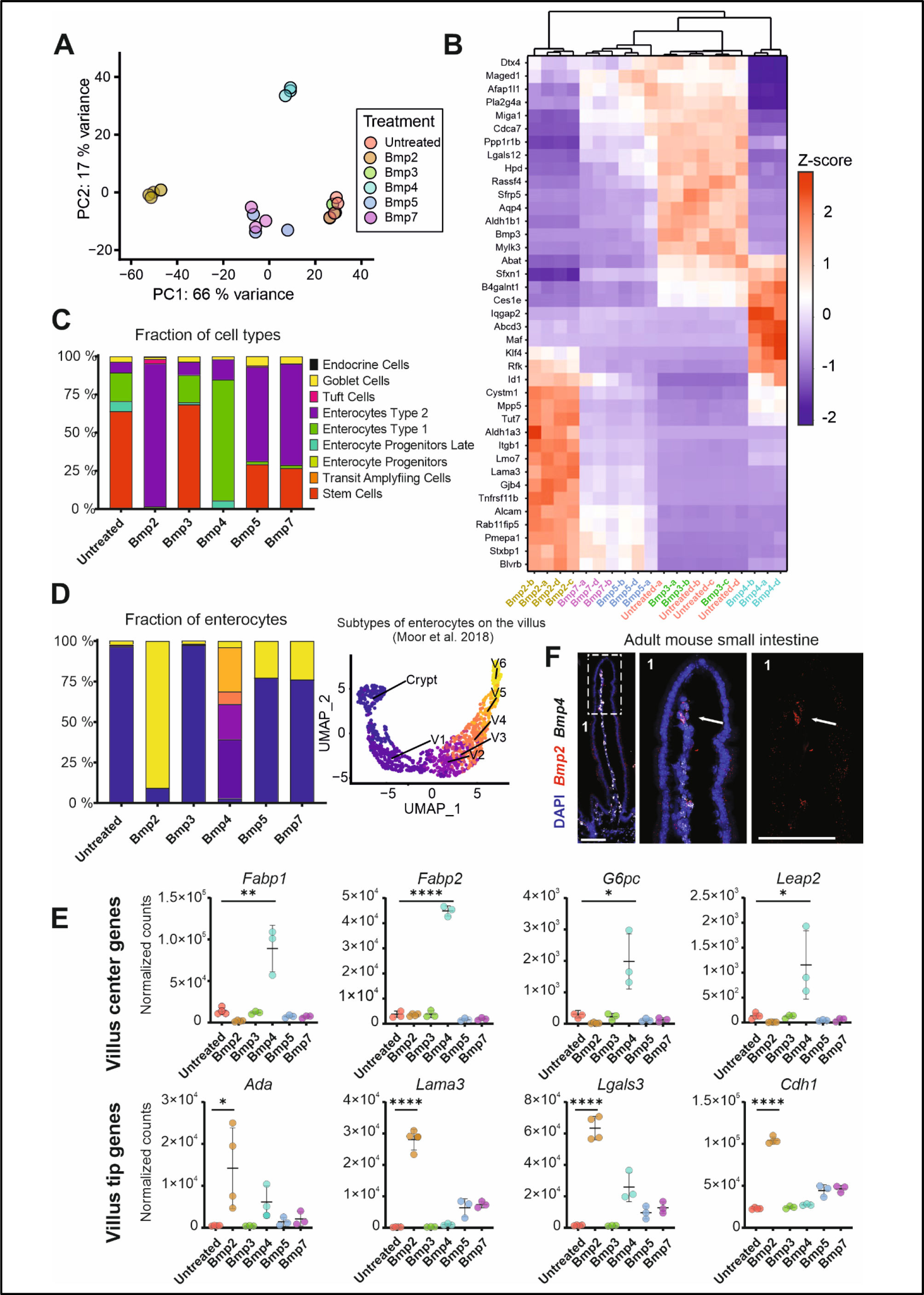
Individual Bmps drive distinct differentiation programs in enterocytes. A Principal component analysis (PCA) of the transcriptomes of intestinal crypts treated with indicated Bmp (500 ng/ml) for 24 hours. PC1 and PC2 explain 66% and 17% of the variance, respectively. (RNAseq samples, n = at least 3 for each Bmp treatment and control). B Heatmap showing changes in expression of indicated genes after 24 hours of indicated Bmp (500 ng/ml) treatment in the intestinal crypts. Hierarchical clustering of Bmp treatments shows differences in gene regulation by Bmp2 vs. Bmp4. (n=3 per each treatment and/or control, top 10 unique genes per each treatment are shown). C Proportional changes of indicated epithelial cell types in the intestinal crypts treated with indicated Bmp (500 ng/ml) for 24 hours. Bmp2 and Bmp4 induce distinct enterocytic programs. (Bar graph depicting deconvoluted RNAseq dataset compared by CellAnneal software to GSE92332 organoids scRNAseq dataset). D Proportional changes of enterocytes subtypes in the intestinal crypts treated with indicated Bmp (500 ng/ml) for 24 hours (left graph). V1 to V6 denote enterocytes according to crypt-villus axes (C = crypt, V1=close to crypt, V6=villus tip) as determined by Moor et al., 2018 and shown on UMAP analysis of 10.5281/zenodo.3403670 dataset (right graph). Colors in the bar graph correspond to enterocyte sub-type from UMAP. (Bar graph depicting deconvoluted RNAseq dataset compared by CellAnneal software to GSM2644349 and GSM2644350 scRNAseq dataset reanalysed by Moor 2018: 10.5281/zenodo.3403670). E Whereas Bmp4 triggers expression of villus-center genes connected to lipid metabolism, Bmp2 preferentially activates the expression of villus tip genes. (Graphs show normalized counts determined by RNAseq, n=3 for each treatment, timepoint: 24 hours, error bars denote sd). F *Bmp2* is expressed by non-epithelial cells at the villus tip, whereas *Bmp*4 is produced along the villus. (single-molecule RNA hybridisation, *Bmp2*-red, *Smad4*-white, DAPI counterstains nuclei, 1 indicate inset shown on the right as magnified, Scale bar = 100 μm)

To identify which cellular programs are governed by individual Bmps, we deconvoluted our RNAseq data using Cellanneal package^18^ with a single cell RNA sequencing (scRNA-seq) dataset of untreated wild type small intestinal organoid as a reference^19^. The deconvolution allowed us to determine cellular composition of the organoids upon Bmp treatments. All the cell types (such as Enterocyte Type 1 and Enterocyte Type 2) were already defined in the reference scRNAseq dataset^19^. In general, the short-term Bmp treatment induced enterocyte differentiation (Fig. 2C). Bmp2 treatment increased the Enterocyte Type 2 signature, corresponding to terminally differentiated enterocytes, whereas Bmp4 promoted differentiation into Enterocyte Type 1, characterized by the expression of metabolic genes. Bmp5 and Bmp7 acted like Bmp2, but they did not reduce the number of IECs so drastically. In addition, as well as triggering an Enterocyte Type 2 program, they stimulated differentiation into goblet cells (Fig. 2C). Since Bmp2 and Bmp4 showed a similar activation of generic targets, but they seemed to act differently, we further focused our comparison mainly on these.

Moor and colleagues assigned enterocytic identities to the cell clusters based on their position along the crypt-villus axis (Crypt, V1-V6). ^5^ Based on this framework Bmp2 induced differentiation into villus-tip enterocytes (V6), whereas Bmp4 triggered differentiation into villus-base and center enterocytes (V1-V4) (Fig. 2D). Many of the Bmp4 induced genes encode for important regulators of lipid metabolism and absorption, such as fatty acid binding proteins *Fabp1* and *Fabp2* (Fig. 2E and Fig. EV 2F), as well as a master regulator of amino acid and lipid uptake *Maf* ^14,15^ (Fig. EV 2E). In contrast to Bmp4, Bmp2 preferentially stimulated the expression of villus-tip genes (V6 fraction) associated with adhesion such as *Laminin3* (*Lama3*), *E-cadherin* (*Cdh1*), *galectin* (*Lgals3*) or modulation of immune system *Adenosine deaminase* (*Ada*) (Fig. 2E). The differential activation of *Fabp1*, *Leap3* and *Lama3* was confirmed by real-time qPCR (Fig. EV 2G).

Taken together these observations suggest that Bmp2 activates villus tip programs and Bmp4 a broader central villus program. To explore this further we looked at the expression pattern of the Bmp2 and Bmp4. mRNA in situ hybridisation (RNAscope) revealed Bmp4 was expressed in non-epithelial cells from the center to the top of the villus. In contrast, but consistent with its proposed action at the villus tip, Bmp2 is produced by a few cells at the tip of the villus (Fig. 2F).

Besides enhanced expression of adhesive molecule, the villus-tip enterocytes specifically express proteins acting as immunomodulators Ada and Nt5e (Cd73). The active transcription of these proteins at the villus tip correlates with the protein expression.^4,5^ Bmp2 treatment specifically induced the transcription of *Ada* and *Nt5e* (*Cd73*) (Fig. 2E and 3A). In contrast to Bmp4, Bmp2 increased the presence of Nt5e (Cd73) positive cells in the intestinal organoids (Fig. 3B,C).

**Figure 3.**
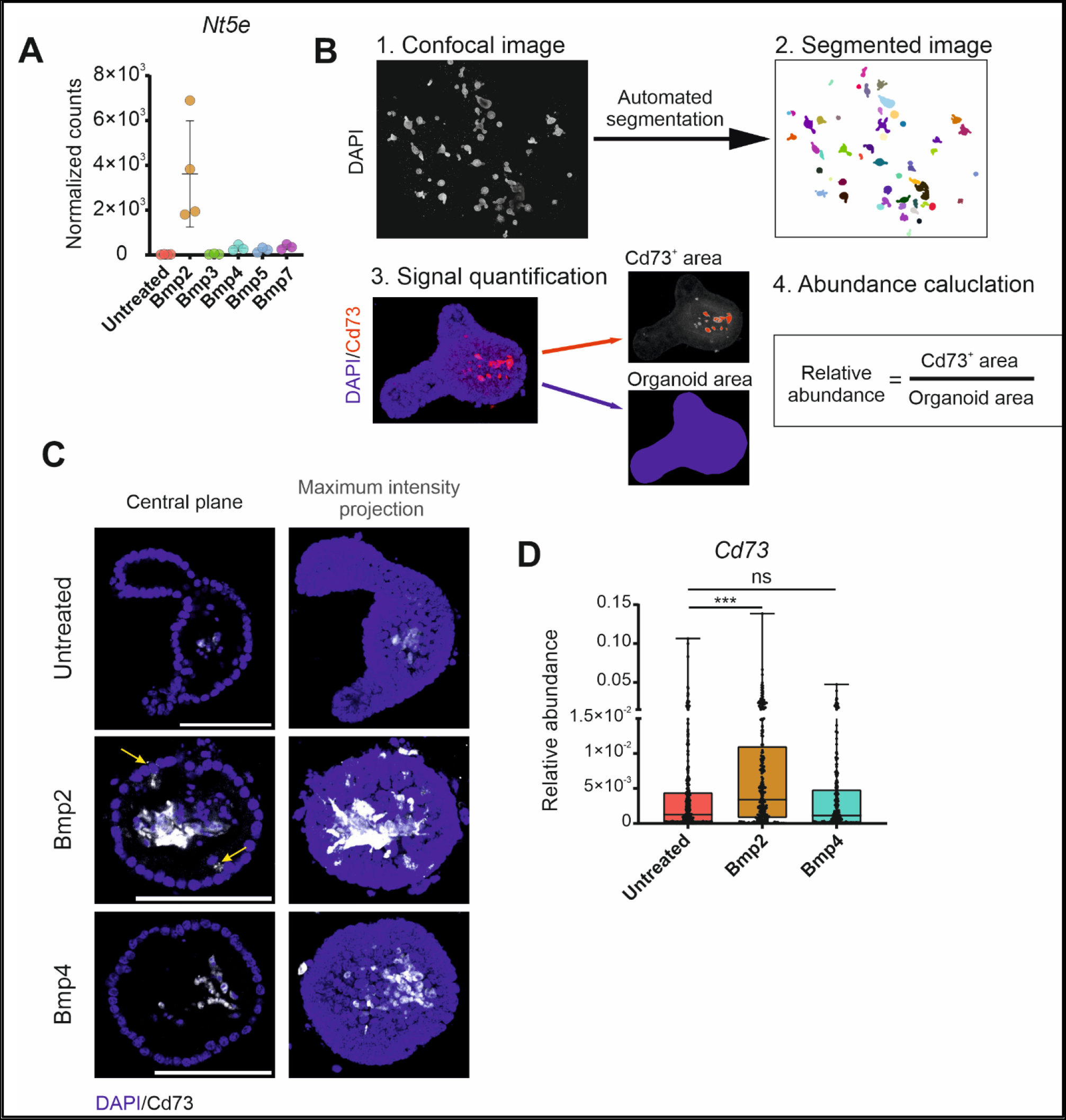
Villus-tip Nt5e-positive cells appear upon Bmp2 treatment. A Bmp2 induce the expression of the villus-tip markers *Nt5e* (*Cd73*). (Graphs show normalized counts determined by RNAseq, n=3 for each treatment, timepoint: 24 hours, error bars denote sd). B Scheme of the experimental pipeline used for immunostaining of Bmp-treated organoids. C Expression of Nt5e/Cd73 (red) upon the Bmp2-treatment in the intestinal organoids. Immunocytochemistry, DAPI (blue) counterstains nuclei, Arrows point to Cd73 positive cells. Scale bar = 100 μm. D Quantification of Nt5e immunostainings determined as the ratio of specific signal to DAPI ratio. (n= at least 100 organoids, One-way ANOVA, graphs show mean ± SD, *** p-value ≤ 0.001).

Altogether, these observations support a positional effect in which subset of villus mesenchymal cells control the villus-tip program in enterocytes by Bmp2.

### Tgfβ promotes villus-tip differentiation of enterocytes similar to Bmp2

Besides Bmps, Pdgfra^high^ villus mesenchymal cells express Tgfβ, (Fig. 1C). Interestingly, in advanced colon cancer, Tgfβ secreted by surrounding mesenchymal cells initiates a Wnt-independency in colon cancer cells^20^. To assess the role of Tgfβ on healthy small intestinal epithelium, we treated intestinal organoids using the same setup as for Bmps. The impact of Tgfβ on the growth and morphology of the organoids was analogous to that of Bmps (Fig. 4A and Fig. EV 3A). Tgfβ seems to activate a program similar to that triggered Bmp2 (respectively Bmp2/5/7) (Fig. 4B,C and Fig. EV 3B-D). Additionally, we saw that Tgfβ treatment reduced expression of villus center genes (Fig. 4C and Fig. EV 3E). We suggest this is because Tgfβ promoted the differentiation of villus-tip enterocytes (Enterocytes Type 2) reduced the number of villus-center enterocytes and consequently the overall levels in these RNAseq experiments (Fig. EV 3D). Of note is the increased fraction of tuft cell (Fig. EV 3D), observed also in the case of Bmp2 treatment (Fig. 2C).

**Figure 4.**
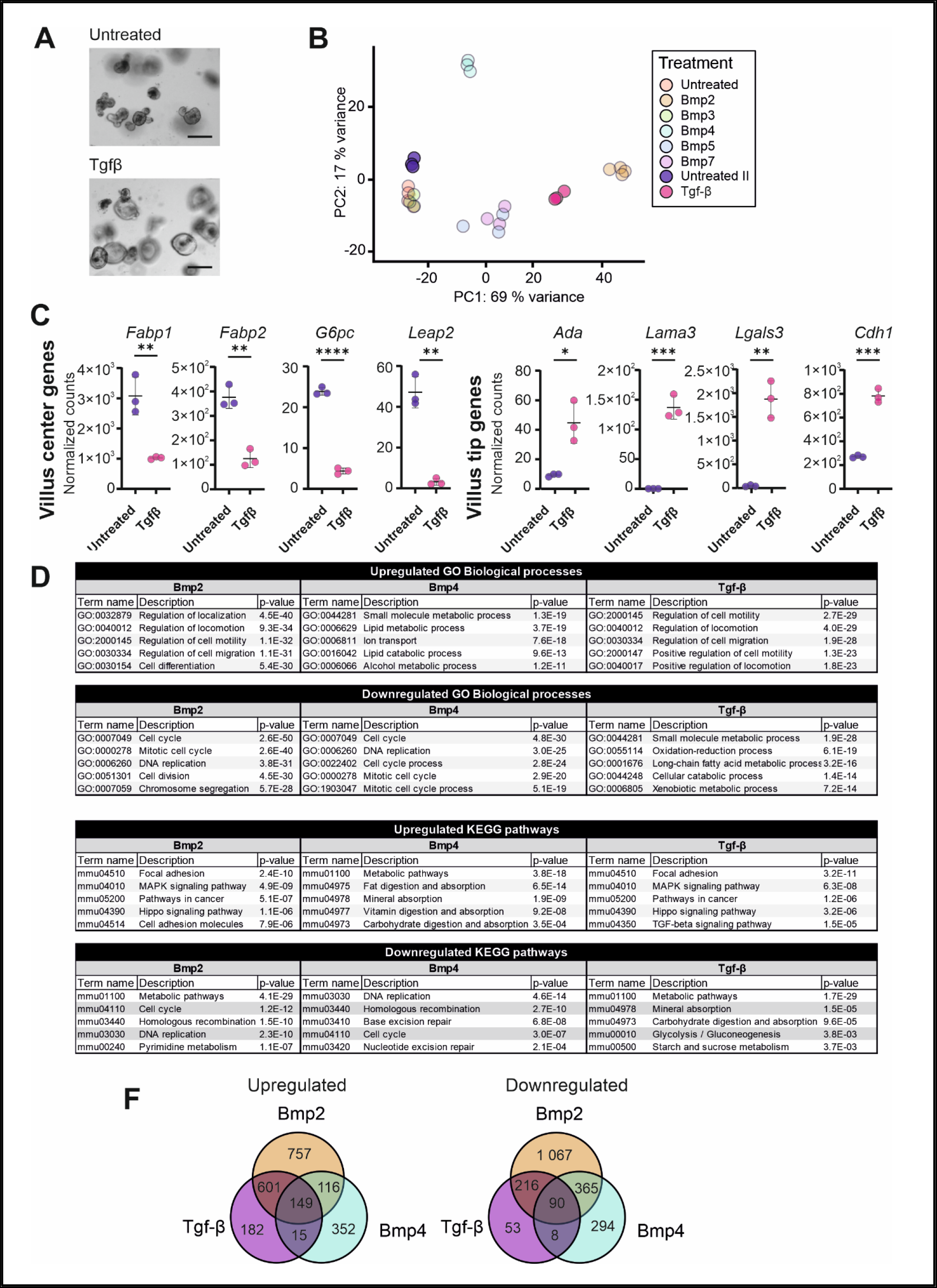
Tgfβ promotes differentiation program similar to Bmp2 and different from Bmp4. A Effect of Tgfβ treatment (after 24 hours, 500ng/ml) on the morphology and growth of freshly isolated intestinal crypts. (Brightfield, Scale bar, 200 μm). B Principal component analysis (PCA) of the transcriptomes of intestinal crypts treated with Tgfβ and indicated Bmp for 24 hours. PC1 and PC2 explain 69% and 17% of the variance, respectively. (RNAseq samples, n=3 for each treatment and control, Tgfβ and Bmp treatment data were normalized and matched based on control/untreated parallels). C Tgfβ induces the expression of villus tip genes and represses villus center genes. (Graphs show normalized counts determined by RNAseq, n=3 for each treatment, timepoint: 24 hours, error bars denote sd). D Relevant upregulated and downregulated gene ontology (GO) terms and KEGG pathways based on differentially expressed genes with │log_2_FC│ ≥ 2, p < 0.05 as calculated using Cytoscape platform (STRING App). E Venn diagrams of intersecting upregulated or downregulated genes │log_2_FC│ ≥ 2, p < 0.05 indicate high overlap between genes regulated by Tgfβ and Bmp2, but low overlap in genes simultaneously regulated by Tgfβ and Bmp4.

The dichotomy of programs regulated by Bmp family members led us to further characterize the impact of Bmp2 and Tgfβ on one branch and Bmp4 on the other, in terms of affected biological processes and altered signalling pathways. As shown before, Bmp4 preferentially upregulated genes connected to metabolism (lipid metabolism) and nutrient absorption, whereas Bmp2/Tgfβ altered cell adhesion, cell migration and locomotion (Fig. 4D). Almost 80% of genes upregulated by Tgfβ are also activated by Bmp2, pointing to their overlapping activity (Fig. 4E). On the contrary, the overlap in upregulated genes between Tgfβ and Bmp4 is lower. Hence GO (Gene ontology) term analysis and KEGG signaling pathway analysis confirmed that whereas Bmp4 governs differentiation of villus-center and enterocytes triggering genes important for lipid metabolism and uptake, Bmp2 and Tgfβ alter the cellular adhesion activating villus-tip programs.

### Villus-tip differentiation driven by Bmp2 depends on canonical BmpRI/Smad4 pathway

Bmp ligands can act via canonical mechanism that starts upon the binding of Bmp to receptors consisting of Type I and Type II forming hetero-tetrameric complexes. Interaction of ligand with the receptor complex initiates a cascade of phosphorylation events ultimately resulting in the phosphorylation of receptor-regulated Smad (R-Smad1/5/8). Phosphorylated R-Smad associates with core Smad4. Consequently, the Smad complex translocates to the nucleus where it triggers the expression of targets genes. However Bmps can also induce target genes expression via various non-canonical, Smad-independent ways^10,11^. To dissect whether Bmp2 activates villus-tip enterocytic programs via the canonical pathway, we administered Bmp type I receptor inhibitor LDN-193189 to wild type mice (Fig. 5A). Short-term repression of Bmp receptor complex (4 days) altered villus-tip differentiation: reduced expression of villus-tip marker genes (Fig. 5B). Of note is the fact that the epithelial morphology and proliferation stayed the same after LDN-193189 treatment (Fig. EV 4B). Intriguingly, villus-center genes were not affected (Fig. EV 4A). Possibly, the Bmp2-driven villus tip program seems to be sensitive to the pathway perturbations, while zonated gene expression within the center of the villus controlled preferentially by Bmp4 might be more resilient. An alternative explanation is the LDN-193189 was not able to penetrate as effectively the central villus region. The requirement of Bmp receptor complex to trigger villus-tip gene expression/program corroborates the results from the organoid-ivMCs co-cultivation model (Fig. EV 1D).

**Figure 5.**
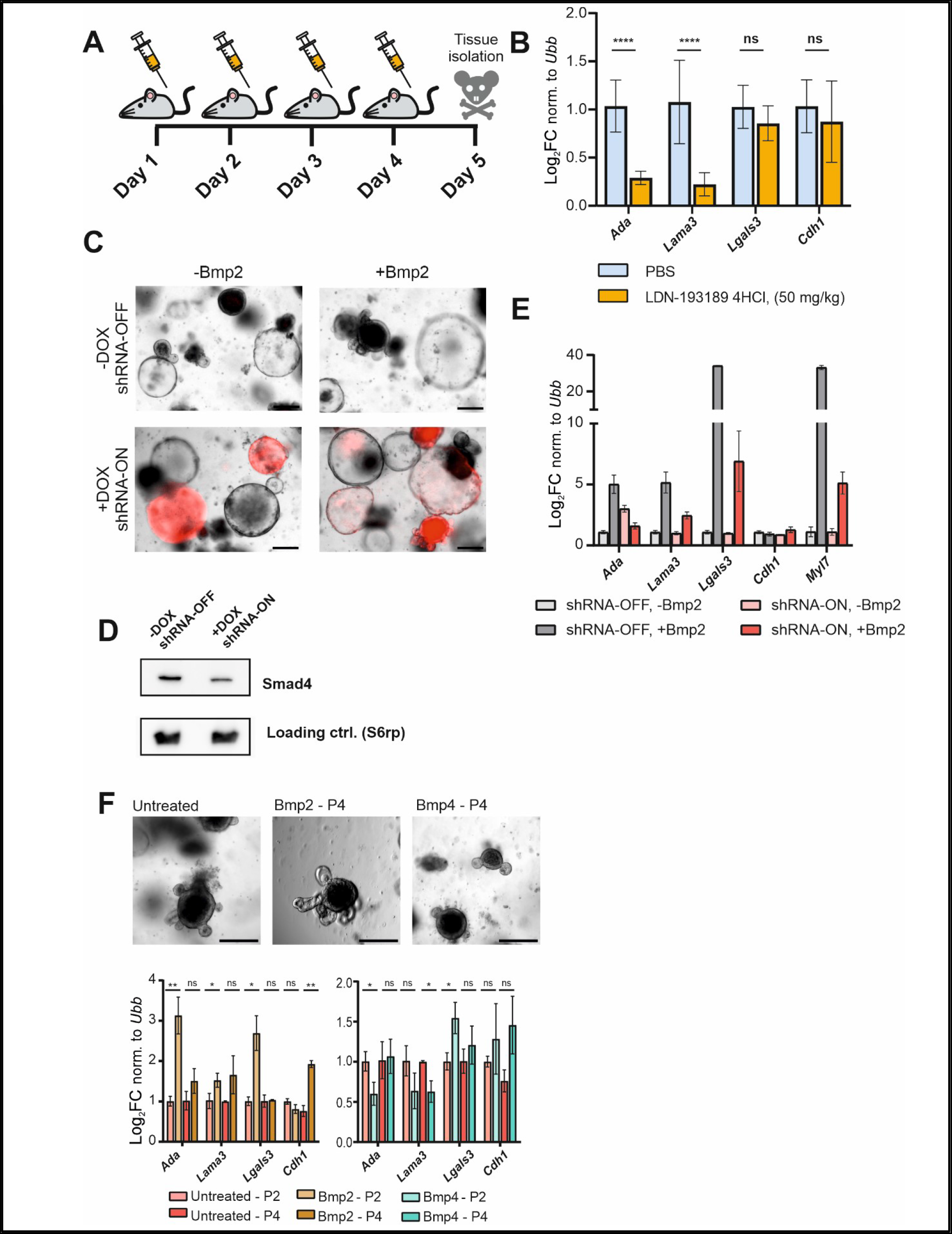
Bmp type I receptor and Smad4 are required to promote Bmp2-driven differentiation programs induced by vMCs. A Scheme of Bmp type I receptor inhibitor LDN-193189 administration. B Four days treatment with LDN-193189 inhibitor reduces the enterocytic villus-tip signature determined by the expression of villus-tip genes genes. (qRT-PCR, expression levels normalized to *Ubiquitin B* (*Ubb*), n=3, error bars show sd, untreated parallel set as 1). C Induction of shRNA against Smad4 upon the doxyciclin (+Dox) treatment (3 days) followed by Bmp2 treatment for additional 24 hours. shRNA activation is visualised by the expression of coupled tdTomato (red). Scale bar = 200 μm D The shRNA against Smad4 (+Dox / shRNA-ON) leads to reduced levels of Smad4. (Immunoblot, after 4 days +/− Dox). E Smad4 knock-down (shRNA-ON) alleviates Bmp2-induced differentiation marked by relative expression of villus-tip genes. (qRT-PCR, expression levels normalized to *Ubiquitin B* (*Ubb*), n=3, error bars show sd, untreated parallel set as 1, treatment as in 4C). F Long term organoid cultivation in the presence of Bmp2 (10ng/ml) enriches the expression of villus-tip genes. (qRT-PCR, expression levels normalized to *Ubiquitin B* (*Ubb*), n=3, error bars show sd, untreated parallel set as 1, P2 – collected 4 days after 2^nd^ passage, P4 - collected 4 days after 4^th^ passage).

To further determine whether Bmp2-induced enterocytic differentiation requires canonical Smad-dependent route, we generated organoids with a doxycyclin-inducible shRNA against Smad4 (shRNA^Smad4^) Despite not all organoids expressed shRNA^Smad4^ as marked by coupled tdTomato, the doxycycline induction reduced expression of Smad4 (Fig. 5C,D). When the organoids with reduced Smad4 were treated with Bmp2 for 24 hours, the activation of villus-tip genes as lower than in wild type organoids (Fig. 5E). The Smad4 depletion also restored proliferation and alleviated IESCs exhaustion caused by Bmp2 treatment (Fig. EV 4C). Importantly, in contrast to freshly isolated crypts used for the organoid experiments so far, shRNA^Smad4^-organoids spent long time in the culture, were transduced and passaged many times. Nevertheless, they still respond to Bmp2 treatment by activating villus-tip program dependent on core Smad4.

### Mimicking the conditions at the villus tip via long term-cultivation in the presence of Bmp2

Long term cultivated intestinal organoids do not completely recapitulate intestinal epithelium: they rarely contain villus-tip cells and most enterocytes have a villus-bottom identity.^19,21,22^ To enrich for villus-tip cells and make organoids more similar to real villi, at least in certain aspects, we tried to establish long term organoid culture using medium contaning low concentrations of Bmp2 (250 ng/ml). Bmp2-treated organoids could survive up to three passages. Using lower Bmp2 concentration (10 ng /ml), the organoids could be propagated in Bmp2-containing medium for minimally 4 weeks (at least 4 passages). Addition of Bmp2 to standard cultivation medium enriched for villus-tip signatures, (Fig. 5F), again underscoring role of Bmp2 as an inducer of villus-tip expression in the enterocytes. Hence, we established a system suitable to test various aspects of villus-tip in the future.

A specific subset of villus-mesenchymal cells - Lgr5 positive mesenchymal cells - are localized in the villus tip. This subset expresses Bmp2, Bmp4 and Wnt5a^16^. Bahan Halpern and collegues showed that the elimination of these cells led to loss of villus-tip identity in enterocytes. ^16^ As we saw with a short-term inhibition of Bmp receptor type I (Fig. 5B), the villus-tip identity was also lost. Our data suggest that Bmp2 could be the main factor controlling this identity. Whether Wnt5a acts synergistically or in parallel to Bmp2 should be investigated in more detail in the future. However, a pilot experiment suggests Bmp2 and Wnt5 can synergize and that Wnt5a can activate villus-tip genes (Fig. EV 4D).

## Discussion

Proper spatial and temporal control of cellular differentiation is an ultimate requirement for the normal intestinal functions. The balance between continual supply of IESCs and differentiation of epithelial progenitors depends on surrounding non-epithelial cells creating the adequate microenvironment. Whereas the role of an intestinal stem cells niche supporting the maintenance of IESCs has been clearly documented in recent years^6,7,9^, the existence and function of a “differentiation niche” remains enigmatic. However, the existence of such a niche is flagged by the presence of villus-localized Pdgfra^high^ mesenchymal cells express mixture of Bmps. Bmp signaling acts as a negative regulator of crypt formation, restricting the stemness^12,23^. Indeed, the epithelial differentiation is not only a default state, that is automatically triggered in the absence of the stem cell programs. The lack of IESCs - promoting signals, does not mean successful differentiation. Bmps actively promote the epithelial differentiation along the crypt-villus axis. Bmps produced by sub-epithelial villus mesenchymal cells contribute to the formation of sequential and zonated gene expression in enterocytes^13^. The Bmp gradient along the villus is classically thought to trigger the sequential gene expression and thus the differentiation^1,2^. But why are there so many distinct Bmps expressed by villus mesenchymal cells if the gradient could simply be formed by a single ligand? What if the identity of the individual Bmps also matters? Our results demonstrate yes, the identity of the individual Bmps matters.

Two highly expressed Bmps are Bmp2 and Bmp4. A combination of Bmp2 and Bmp4 was shown to push the differentiation of enterocytes by activating genes involved in lipid metabolism and uptake^13^. Two studies highlighted c-Maf as a key regulator of this metabolic differentiation^14,15^. One study showed Bmp2 induces expression of c-Maf, while the other described Bmp4 as the trigger (both *in vitro*)^14,15^. We also found that, both Bmp2 and Bmp4 can induce *Maf* but that Bmp4 was much more potent (Fig. EV 2E). Crucially, however, our transcriptome analysis suggests that Bmp2 and Bmp4 trigger distinct programs Whereas Bmp4 governs the lipid metabolism and uptake, Bmp2 regulates the cellular locomotion, adhesion and expression of immunomodulators (Fig.2E, Fig.3A). The activation of distinct programs can be connected to expression pattern of Bmps. Whereas Bmp4 is expressed along the villus, Bmp2 is produced in few cells at the very tip of the villus (Fig.2F). The pattern of expression corresponds to enterocytic zonation – enterocytes in the middle to the top of the villus express metabolic genes, whereas villus-tip enterocytes are characterized by changed adhesion and production of immunomodulators^4,5^ (Fig.2D,E).

The role of Bmp2 in priming villus-tip identity goes in the line with the observation that rare Lgr5-positive cells localised at the villus tip promote the terminal differentiation of enterocytes^16^ Besides Bmp2 and Bmp4, Lgr5 cells also secrete Wnt5a. Our pilot experiments suggest the potential of Wnt5a to induce villus-tip features in parallel to Bmp2. Importantly, there might be a synergy between Bmp2 and Wnt5a in this process (Fig. EV 4D). The fact, that Wnt5a could not induce the expression of villus-center genes (lipid metabolism and uptake)^13^ may point to specific synergism between Bmp2 and Wnt5a, since Bmp2 is not expressed at the center of the villus. Nevertheless, further research (beyond this work) is required to confirm or disprove this phenomenon.

Villus-tip epithelial identity is triggered by Bmp2 in canonical, Smad-dependent way (Fig. 5 B,E). The enterocytes quickly move upwards along the villus. Thus, the time dictates how long the distinct Bmp signalling can act. Whereas the short-term exposure drives preferentially enterocytic programs (Fig. 2C,D), longer exposure additionally drives the differentiation into goblet cells (Fig. 1F), as also shown recently^13^.

Bmp5 and Bmp7 are the other ligands expressed by Pdgfra^high^ villus mesenchymal cells. They tiggered a program similar to that elicited by Bmp2 (Fig. 2D, E). It goes in the line with what is known about Bmps homo- or heterodimerization. Bmp5 heterodimerizes just with Bmp2, Bmp7 can form heterodimers with both Bmp2 and Bmp4. On the other hand, Bmp2 does not heterodimerize with Bmp4^10,11,17^. This may indirectly support the distinct roles of Bmp2 vs. Bmp4. Regulation of adhesive properties can also be modulated by Tgfβ, acting similarly as Bmp2 on healthy organoids. Of note also Tgfβ elicited a differentiation program similar to that of Bmp2 (Fig. 3D,E). Tgfβ seems to act mainly as immunoregulator in the healthy intestine^24,25^, but older reports indicate it can affect the intestinal epithelium, consistent with our observations. Tgfβ enhances the cellular adhesion of intestinal epithelial cell line grown as monlayer *in vitro*^26^. In the colon, Tgfβ promotes the regeneration of the intestinal epithelium after the mechanical damage^27^. The mechanisms at least partially depends on affecting the focal adhesion that limits the proliferation, finally facilitating the reappearance of new colonic crypts.^27^ Hence, Tgfβ may act as an alternative regulator of cellular adhesion, activating the gene signature present in villus-tip cells.

Another important aspect of our work is the experimental design and establishment of a new organoid culture system. Since intestinal organoids cultivated long-term under the standard conditions do not recapitulate the composition of differentiated enterocytes^13,19,21^, we used freshly isolated crypts to keep the responsiveness to differentiating stimuli.

Taking together, we showed that individual Bmp ligands secreted by sub-epithelial mesenchymal cells can induce distinct differentiation programs, especially in enterocytes. Whereas Bmp4 activates lipid metabolism and uptake in the center of the villus, Bmp2 modulates properties of villus-tip enterocytes, governing their terminal differentiation. The specific role of Bmp2 can be connected to its expression at the villus tip. Hence the division of labor in enterocytes may at least partially mirror the specific action and expression of particular Bmps.

## Limitation of the study

The key experiments are based on freshly established intestinal organoids the situation in vivo may be more complex. However, the in vivo the villus-tip specific expression of Bmp2 correlates with the gene signature of villus-tip enterocytes that is achieved also by Bmp2 treatment in organoids. Another qualifier that needs to be noted is that the study focuses to proximal small intestine. Finally, the work focuses to describe the impact of individual roles of Bmps on the gene expression of the epithelial cells. Hence the mechanism, how distinct expression of Bmps is controlled is not the part of this work.

## Data availability

Data generated in this study are deposited in GEO with the accession numbers: GSE216699 (https://www.ncbi.nlm.nih.gov/geo/query/acc.cgi?acc=GSE216699, reviewer’s token: ejsraqkutxezlgl). Source data are provided with this paper. The remaining data are available within the Article, Supplementary Information or available from the authors upon the request.

## Acknowledgements

O. Stepanek kindly shared Actin-Cre animals with us. For the technical help we thank Genomic and Bioinformatic Facility of IMG, Prague directed by M. Kolar. This work was supported by Czech Science Foundation grant 21-26025S and The project National Institute for Cancer Research (Programme EXCELES, ID Project No. LX22NPO5102) - Funded by the European Union - Next Generation EU.

## Author’s contribution

L.B. designed and performed experiments, collected and analyzed the data, conducted the bioinformatic analysis. T.V. initiated and concepualized the research, designed the expriments, discussed and interpreted the data and supported the research. The following co-athors assisted with experiments: H.F. (RNAScope), M.S and D.H. (shRNA^Smad4^ organoids), M.D.B. (RNAScope). J.K. helped with the initial bioinformatic analysis. L.B. and T.V. wrote the manuscript. H.F., M.D.B., G.H., V.K. and K.B. critically commented the manuscript.

## Methods

### Mice

Wild-type (*C57BL/6*) mice were obtained from own breeding colony housed at the Institute of Molecular Genetics, Prague, Czech Republic. Pdgfra^H2BeGFP (28)^ was purchased from Jackson Laboratories, United States of America (Stock number:007669). Td Tomato-positive crypts were isolated from Actin-Cre (Gt(ROSA)26Sor(ACTB-Cre,-EGFP)) crossed to Ai14 reporter strain (B6;129S6-Gt(ROSA)26Sortm14(CAG-tdTomato)Hze/J)). Mice were 10-15 weeks old at the time of treatments and crypt isolations, both sexes were used.

To block Bmp type I receptors wild-type animals were intraperitonelly injected with LDN-193189-HCl dissolved in PBS (10 mg/kg, in 100 μl volume, one injection per day) or vehicle (PBS) for 4 consecutive days. The mice were sacrificed 24 h after the last injection.

Housing of mice and in vivo experiments were performed in compliance with the European Communities Council Directive of 24 November 1986 (86/609/EEC) and national and institutional guidelines. Animal care and experimental procedures were approved by the Animal Care Committee of the Institute of Molecular Genetics (no. 2482/2021).

### Mesenchymal cells isolation and immortalisation

Mesenchymal cells from duodenum of Pdgfra^H2BeGFP^ mouse were isolated as described previously^8,9^. The duodenal tissue was harvested, opened longitudinaly and rinsed. The tissue was gently rocked for 30 minitues at room temperature in Gentle Cell Dissociation Reagent to remove epithelial cell. After the detachment of epithelium remaining pieces were digested for 35 minutes in DMEM supplelemented with 1 mg/ml collagenase D and 0,3 mg/ml dispase in thermomixer at 37°C with gentle shaking (60 rpm in horizonally oriented 50ml Falcon tube). Afterwards cells were filtered through 70μm cell strainer and seeded. 24 h after seeding non-adherent cells were washed out. The cells were cultivated in MesenCult medium. The medium was changed 3 times a week. The mesenchymal cells were immortalised by transduction of (HPV16)-E6 gene using retroviral particles. For the retroviral production PlatinumE cells were transfected with pMXs-EF1-Puro retroviral construct containing (HPV16)-E6 insert re-cloned from p1322HPV-16-E6 vector. Lipofectamine 3000 and OptiMEM medium were used according to manufacturers protocol. Transfection medium was changed for cultivation medium 6 h after transfection. Viral particles were produced for 72 h. Retroviral partciles were concentrated overnight at 4 °C by RetroX concentrator and used for infection of mesenchymal cells by spinoculation (30 °C, 450 g,1 h). 72 hours after the spinoculation the medium with viral particles was changed and cells were cells were selected with puromycin (1 μg/ml) and further expanded. The immortalized cells were sorted based on internal eGFP fluorescence intensity to eGFP^low^ eGFP^high^ and further separatelly expanded. Occassionally, the cells were retreated with puromycin. The cells were passaged with TripleExpres solution.

### Crypt isolation and culture

Crypts from proximal small intestine were isolated and cultured as described^29,30^. The dissected intestine was flushed out and opend longitudinally. Villi were shaved off by cover glass. After repeted washing in ice-cold PBS the fragments were gently rocked for 30 min at 4 °C in 5mM EDTA in PBS. Crypts were released by gentle hand shaking for 30 s into ice cold PBS, filtered through 70 μm cell strainer and seeded into Matrigel domes on 24 well plates (3 domes/25 μl per well) and cultivated in complete organoid medium supplemented with mEGF (50 ng/ml), mNoggin (100 ng/ml) and 10 % of Rspondin1 conditioned medium prepared using Cultrex HA-R-Spondin1-Fc 293T cell line according to cell line datasheet– ENR medium. Individual recombinant Bmp proteins or Tgfβ (500 ng/ml) were added to ENR medium at the time of seeding and crypts were cultivatited with Bmps or Tgfβ for 96 hours (with one medium change after 48 hours) for initial experiments shown in Fig.1, Fig.S1 and Fig.S3A). For EdU incroporation assay the medium was changed daily.

For RNAseq and subsequent confirmative quantitative real time PCR experiments, the cypts were intially seeded in ENR medium (without any Bmp/Tgfβ). After 48 h the debris was removed: old medium was removed and matrigel domes containing organoids were broken with 1 ml plastic tip with cut-off tip and transferred into 15 ml tube containing 10 ml of ice-cold PBS. The crypts were incubated 10 min on ice, sedimented and then upper 8 ml of PBS containing cell debris was sucked out and replaced by fresh PBS. Organoids were centrifugated (300 g, 4 °C, 5 min) and put into fresh matrigel domes. Recombinant Bmps or Tgfβ (500 ng/ml) were added for 24 h. For combined Bmp2+Bmp4 treatment the total concentration (500 ng/ml) was kept – i.e. 250 ng/ml of individulal Bmp.

When crypts were cultivated for Western blotting the debris removal step was omitted and organoids were kept in recombinant Bmps for 48 h without medium change.

For the long-term villus-tip enriched culture ENRB medium was used, coposed of ENR supplemented with Bmp2 or Bmp4 (10 ng/ml or 250 ng/ml). The organoids were passaged approximately once in a week.

### EdU staining and image quantification

Duodenal crypts were isolated as mentioned above and seeded on glass-bottomed 8-well chamber (10 μl matrigel drops) for 5 days in ENR medium containing individual Bmps (500 ng/ml). After 96 hours, they were incubated in medium containing 10 μM EdU for 45 min and stained with EdU Cell Proliferation Kit according to manufacturer’s protocol. Afterwards the organoids were counterstained with DAPI. Z-stack confocal images of individual organoids were taken using Dragonfly 503 spinning disc microscope (Andor). EdU and DAPI signal volumes were computed using Fiji software. Finally, (EdU overlapping with DAPI)/DAPI ratio (i.e. percentage of proliferating cells) was calculated. For automated quantification, we used our macro written by *Light Microscopy* core facility at Institute of Molecular Genetics of the Czech Academy of Sciences.

### Co-cultivation of vMCs with intestinal crypts

Villus mesenchymal cells (vMCs) corresponding to immortalized eGFP^high^ cells were seeded on 96 well plate two days before the addition of organoid fragments. TdTomato-positive crypts were isolated and cultivated as described above in advance to co-cultivation start. For the co-cultivation experiment organoids were mechanically dissociated by pipeting. Organoid chunks were mixed in 10 % Matrigel dissolved in ENR organoid medium. 20 μl of crypt/matrigel/ERN medium suspension containing fragments from roughly 10-20 organoid fragments were added to vMCs (70-80 % confluent at the time of mixing) or to empty wells. The plate was incubated 10 min at 37 °C and afterwards gently covered with 100 μl complete ENR organoid medium optionally containing LDN-193189 (1 μM). The co-cultures were analyzed after 3 days.

### Organoid immunostaining, imaging and image analysis

Organoids were generated from isolated crypts of the murine small intestine as done previously^31^. Organoids were kept in IntestiCult Organoid Growth Medium (STEMCELL Technologies) with 100μg/ml Penicillin-Streptomycin for amplification and maintenance. Mechanically split organoids were plated in a 96-well plate and treated with 500 ng/ml BMP2 or 500 ng/ml BMP4 in ENR medium, for 48 hours until the fixation. The procedure for immunostaining was performed as descripted before^31^. Primary antibody of CD73 (BioLegend), 1:200) and secondary antibody (Thermo Fisher Scientific, 1:2000) were diluted in blocking buffer and applied as indicated in Yang et al.^31^ Cell nuclei were stained with 20 μg/ml DAPI Invitrogen) in PBS for 10 min at room temperature.

High-throughput imaging was done with an automated spinning disk microscope from Yokogawa (CellVoyager 7000S), with an enhanced CSU-W1 spinning disk (Microlens-enhanced dual Nipkow disk confocal scanner), a 40x (NA = 0.95) Olympus objective, and a Neo sCMOS camera (Andor, 2,560 × 2,160 pixels). For imaging, an intelligent imaging approach was used in the Yokogawa CV7000 (Search First module of Wako software 2.0) as described before^31^. Z-planes spanning a range up to 80 μm and 2 μm z-steps were acquired. Organoid segmentation in maximum intensity projections (MIPs) was adapted from before^31^. For each acquired confocal z-stack field, MIPs were generated. All MIP fields of a well were stitched together to obtain MIP well overviews for each channel. The high resolution well overviews were used for organoid segmentation and feature extraction. Organoid segmentation based on DAPI channel were generated by a full convolutional neural network (FCN) as in Yang et al.^31^ Artifacts of segmentations were identified by setting size-threshold and visual inspection, annotated in Fiji (Version 2.9.0) and removed by a customized python script. From the segmented MIPs, we calculate the size of each individual organoid from label masks. Using the individual organoid masks from the MIP, we crop for each organoid the signals of all recorded channels. Uniformed threshold for CD73 intensity was applied to all samples to generate the segmentation of CD73^+^ regions. Then, the CD73 ratio of each individual organoid was calculated by the size of CD73^+^ region dividing the size of organoid label mask (see Fig. 3 B).

### Smad4 shRNA organoids line preparation and experiment

Lentiviral construct containing doxycycline inducible shRNA was generated by inserting oligos targeting Smad4 (see the Key Resource table for sequences) into pTripz vector. Viral particles were prepared according to pTRIPZ Technical manual in HEK 293 FT cell line using Lipofectamine 2000 for the transfection of lentiviral vectors. After 72 hours, lentiviral suspension was collected and concentrated overnight using Lenti-X concentrator. Small intestinal organoids were pretreated with WntSur 0,5uM, Nicotinamide 10mM, Y-27632 10μM for 96 hours before the lentiviral infection. The organoids were disintegrated to single cell suspension using TripleExpress reagent. The cells were spinoculated with concentrated viral suspension (32°C, 600g, 1 hour) and kept in 37 °C for another 5 hours. Afterwards they were embedded into matrigel and kept in ENR containing WntSur 0,5uM, Nicotinamide 10mM, Y-27632 10μM. Futher the transduced organoids were selected using puromycin 2 μg/ml.

Before sorting, organoids were pretreated with doxycycline 1 μg/ml. At the day of sort, organoids were disintegrated to single cell suspension using TripleExpress reagent. Cells with the highest RFP signal were sorted into ENR containing WntSur 0,5uM, Nicotinamide 10mM, Y-27632 10μM and 10 % matrigel, centrifugated and embedded into matrigel and medium containing WntSur 0,5uM, Nicotinamide 10mM, Y-27632 10μM. The sorting step was repeated twice. Finally, WntSur 0,5uM, Nicotinamide 10mM, Y-27632 10μM were gradually removed. The oragnoids were occasionally reselected with 2 μg/ml puromycin For the experiment organoids were passaged and treated +/− doxycyclin (1 μg/ml) for 3 days. Afterward Bmp2 (+/− doxycyclin) or vehicle (+/− doxycyclin) were added and organoids were cultured for additional 24 hrs. Then they were photographed and harvested. Note: Despite repeated selection and sorting not all organoids expressed shRNA^Smad4^ as marked by coupled tdTomato upon Dox treatment.

### Histology, immunohistochemistry, microscopy

Dissected duodenum was flushed out by ice-cold PBS, then cut into 1 cm pices and fixed in 4%PFA overnight at 4°C. Afterwards tissues were repeatedly washed with PBS, dehydrated, embedded in parrafin and cut 5 μm sections. Deparaffinized tissue sections were subjected to antigen retrieval in 2.4 mM sodium citrate and 1.6 mM citric acid, pH 6, for 25 min in a steamer. Sections were washed with PBST (0.1% Tween-20 in PBS) and blocked for 30 min at RT in blocking buffer (5% BSA, 5% heat-inactivated normal goat serum in PBST). Following overnight incubation at 4°C with primary antibody (1:100, in blocking buffer), sections were washed in PBST and incubated with secondary antibody (1:400, in blocking buffer) for 1 hr at room temperature. The antibodies are indicated in Key Resource Table. Nuclei were stained with DAPI (1:1000) in blocking solution for 5 min at RT. Sections were imaged on DM6000 (Leica).

### Quantitative real-time PCR

For cultured crypts, matrigel was washed out by ice-cold PBS and cells peletted. The intestinal epithelium from animals was isolated as descibed^32^. Total RNA from crypts or intestinal epithelium was extracted using RNeasy Micro Kit according to manufacturer’s protocol including DNAseI treatment. cDNA was synthetized with RevertAid First Strand cDNA Synthesis Kit. qRT-PCR was performed in three replicates. Fold change values were counted using ΔΔC_T_ method.

### In situ mRNA hybridisation

mRNAs were detected and visualized using RNAscope method (Advanced Cell Diagnostics, Germany) on small intestine tissue sections according to manufacturer’s protocol (RNAscope Fluorescent Multiplex Assay). Probe sets for *Bmp2* and *Bmp4* were designed by Advanced Cell Diagnostics. Images were taken by Leica SP8 laser scannicg confocal microscope.

### Western blotting

Cultivated crypts were harvested and treated with Recovery Solution for 30 min on ice to remove residual matrigel. Afterwards they were pelleted and lysed in buffer containing: 50 mM HEPES/KOH pH 7.4, 1% Triton X-100, 50 mM NaF, 5 mM Na_2_H_2_P_2_O_7_, 400 mM NaCl, 40 mM β-glycerolphosphate, 12.5 mM EGTA pH 8, 1 mM Na_3_VO_4_, 1 mM benzamidin, 0.5 mM PMSF, 1.5 mM MgCl_2_, leupeptin 0.04 μl/ml, pepstatin 0.04 μl/ml, antipain 0.04 μl/ml.Total concentration of proteins was determined using standard Bradford assay.

Proteins were separated on 15% acrylamide-Tris-BIS-0.1% SDS gel, 6,16 % acrylamide-Tris-BIS gel was used as stacking. Afterwards proteins were transferred to nitroceluulose membrane using semi-dry Blotter BioRad (2,5 mA/cm2, 10 W, 15 V, 55 min). All antibodies were diluted according to manufacturer’s recommendations. Antibodies were visualised with chemiluminiscent substrate SuperSignal West Pico PLUS. Used antibodies are indicated in Key Resource Table.

### RNA sequencing

For control samples (untreated samples, at least n=3 independent experiments, each consisted of pooled technical replicates) and experimental samples (individual Bmp or Tgfβ treatmet for 24 hours, for each Bmp, n=3 independent experiments, each consisted of pooled technical replicates) total RNA was isolated with the same procedure as for qRT-PCR. Sequencing libraries were prepared from total RNA using the KAPA mRNA Hyperprep Kit, followed by size distribution analysis in the Agilent 2100 Bioanalyzer using the High Sensitivity DNA Kit (Agilent). Libraries were sequenced in two runs of the Illumina NextSeq 500 instrument (Illumina, USA) using a 75nt single-end configuration. The complete processing of isolated RNA was done by Genomics and Bioinformatics core facility at Institute of Molecular Genetics of the Czech Academy of Sciences.

### Computational RNA sequncing analysis

The nf-core/rnaseq bioinformatics pipeline version 3.5 ^33^ was used for subsequent read processing. Individual steps included removal of sequencing adapters and low-quality reads with Trim Galore! (http://www.bioinformatics.babraham.ac.uk/projects/trim_galore) and quantification of gene expression with Salmon ^34^ using GRCm39 reference^35^. Estimated expression per gene served as input for differential expression analysis using the DESeq2 R Bioconductor package^36^. Genes exhibiting a minimum absolute log2 fold change of 1 (|log_2_ FC| ≥ 1) and statistical significance (adjusted p-value < 0.05) between the compared sample groups were considered differentially expressed. The initial mapping and analysis of sequencig data was done by Genomics and Bioinformatics core facility at Institute of Molecular Genetics of the Czech Academy of Sciences.

#### Heatmap

For each treatment top 10 differentially expressed genes were selected. Z-score was calculated using standard formula z = (x – μ) / σ. The heatmap was generated with heatmaply package in R.

#### Deconvolution of RNAseq datasets

The percentage values charts were created by using Cellanneal softwere with default deconvolution values kept. Normalized RNA counts from RNAseq experiments were used. Pseudobulked datasets from Haber et al., 2017^19^ (GSE92332) and Moor et al., 2018^5^ https://zenodo.org/record/3403670) were used as reference datasets for different visualisations. The datasets were preprocessed using standard Seurat forkwlow. Biological replicates were averaged. The code used for preprocessing of raw scRNA-seq count matrices is available.

## References

1. Spit, M., Koo, B.-K. & Maurice, M. M. Tales from the crypt: intestinal niche signals in tissue renewal, plasticity and cancer. Open Biol. 8, 180120 (2018).

2. Gehart, H. & Clevers, H. Tales from the crypt: new insights into intestinal stem cells. Nat Rev Gastroenterol Hepatol 16, 19–34 (2019).

3. Schwanhäusser, B. et al. Global quantification of mammalian gene expression control. Nature 473, 337–342 (2011).

4. Harnik, Y. et al. Spatial discordances between mRNAs and proteins in the intestinal epithelium. Nat Metab 3, 1680–1693 (2021).

5. Moor, A. E. et al. Spatial Reconstruction of Single Enterocytes Uncovers Broad Zonation along the Intestinal Villus Axis. Cell 175, 1156–1167.e15 (2018).

6. Greicius, G. et al. *PDGFRα ^+^* pericryptal stromal cells are the critical source of Wnts and RSPO3 for murine intestinal stem cells in vivo. Proc Natl Acad Sci USA 115, E3173–E3181 (2018).

7. McCarthy, N. et al. Distinct Mesenchymal Cell Populations Generate the Essential Intestinal BMP Signaling Gradient. Cell Stem Cell S1934590920300084 (2020) doi:10.1016/j.stem.2020.01.008.

8. Brügger, M. D., Valenta, T., Fazilaty, H., Hausmann, G. & Basler, K. Distinct populations of crypt-associated fibroblasts act as signaling hubs to control colon homeostasis. PLoS Biol 18, e3001032 (2020).

9. Degirmenci, B., Valenta, T., Dimitrieva, S., Hausmann, G. & Basler, K. GLI1-expressing mesenchymal cells form the essential Wnt-secreting niche for colon stem cells. Nature 558, 449–453 (2018).

10. Wang, R. N. et al. Bone Morphogenetic Protein (BMP) signaling in development and human diseases. Genes & Diseases 1, 87–105 (2014).

11. Salazar, V. S., Gamer, L. W. & Rosen, V. BMP signalling in skeletal development, disease and repair. Nat Rev Endocrinol 12, 203–221 (2016).

12. Qi, Z. et al. BMP restricts stemness of intestinal Lgr5+ stem cells by directly suppressing their signature genes. Nat Commun 8, 13824 (2017).

13. Beumer, J. et al. BMP gradient along the intestinal villus axis controls zonated enterocyte and goblet cell states. Cell Reports 38, 110438 (2022).

14. Cosovanu, C. et al. Intestinal epithelial c-Maf expression determines enterocyte differentiation and nutrient uptake in mice. Journal of Experimental Medicine 219, e20220233 (2022).

15. González-Loyola, A. et al. c-MAF coordinates enterocyte zonation and nutrient uptake transcriptional programs. Journal of Experimental Medicine 219, e20212418 (2022).

16. Bahar Halpern, K. et al. Lgr5+ telocytes are a signaling source at the intestinal villus tip. Nat Commun 11, 1936 (2020).

17. Bragdon, B. et al. Bone Morphogenetic Proteins: A critical review. Cellular Signalling 23, 609–620 (2011).

18. Buchauer, L. & Itzkovitz, S. cellanneal: A User-Friendly Deconvolution Software for Omics Data. (2021) doi:10.48550/ARXIV.2110.08209.

19. Haber, A. L. et al. A single-cell survey of the small intestinal epithelium. Nature 551, 333–339 (2017).

20. Han, T. et al. Lineage reversion drives WNT independence in intestinal cancer. http://biorxiv.org/lookup/doi/10.1101/2020.01.22.914689 (2020) doi:10.1101/2020.01.22.914689.

21. Grün, D. et al. Single-cell messenger RNA sequencing reveals rare intestinal cell types. Nature 525, 251–255 (2015).

22. Fujii, M. et al. Human Intestinal Organoids Maintain Self-Renewal Capacity and Cellular Diversity in Niche-Inspired Culture Condition. Cell Stem Cell 23, 787–793.e6 (2018).

23. Haramis, A.-P. G. et al. De Novo Crypt Formation and Juvenile Polyposis on BMP Inhibition in Mouse Intestine. Science 303, 1684–1686 (2004).

24. Bauché, D. & Marie, J. C. Transforming growth factor β: a master regulator of the gut microbiota and immune cell interactions. Clin Trans Immunol 6, e136 (2017).

25. Stolfi, C., Troncone, E., Marafini, I. & Monteleone, G. Role of TGF-Beta and Smad7 in Gut Inflammation, Fibrosis and Cancer. Biomolecules 11, 17 (2020).

26. Howe, K. L., Reardon, C., Wang, A., Nazli, A. & McKay, D. M. Transforming Growth Factor-β Regulation of Epithelial Tight Junction Proteins Enhances Barrier Function and Blocks Enterohemorrhagic Escherichia coli O157:H7-Induced Increased Permeability. The American Journal of Pathology 167, 1587–1597 (2005).

27. Miyoshi, H., Ajima, R., Luo, C. T., Yamaguchi, T. P. & Stappenbeck, T. S. Wnt5a Potentiates TGF-β Signaling to Promote Colonic Crypt Regeneration After Tissue Injury. Science 338, 108–113 (2012).

28. Hamilton, T. G., Klinghoffer, R. A., Corrin, P. D. & Soriano, P. Evolutionary Divergence of Platelet-Derived Growth Factor Alpha Receptor Signaling Mechanisms. Mol Cell Biol 23, 4013–4025 (2003).

29. Sato, T. et al. Single Lgr5 stem cells build crypt-villus structures in vitro without a mesenchymal niche. Nature 459, 262–265 (2009).

30. Sato, T. et al. Paneth cells constitute the niche for Lgr5 stem cells in intestinal crypts. Nature 469, 415–418 (2011).

31. Yang, Q. et al. Cell fate coordinates mechano-osmotic forces in intestinal crypt formation. Nat Cell Biol 23, 733–744 (2021).

32. Gracz, A. D., Puthoff, B. J. & Magness, S. T. Identification, Isolation, and Culture of Intestinal Epithelial Stem Cells from Murine Intestine. in Somatic Stem Cells (ed. Singh, S. R.) vol. 879 89–107 (Humana Press, 2012).

33. Ewels, P. A. et al. The nf-core framework for community-curated bioinformatics pipelines. Nat Biotechnol 38, 276–278 (2020).

34. Patro, R., Duggal, G., Love, M. I., Irizarry, R. A. & Kingsford, C. Salmon provides fast and bias-aware quantification of transcript expression. Nat Methods 14, 417–419 (2017).

35. Howe, K. L. et al. Ensembl 2021. Nucleic Acids Research 49, D884–D891 (2021).

36. Love, M. I., Huber, W. & Anders, S. Moderated estimation of fold change and dispersion for RNA-seq data with DESeq2. Genome Biol 15, 550 (2014).

