## Supplementary Figures for "Terminal differentiation of villus-tip enterocytes is governed by distinct members of Tgfβ superfamily"

*Figure EV 1*

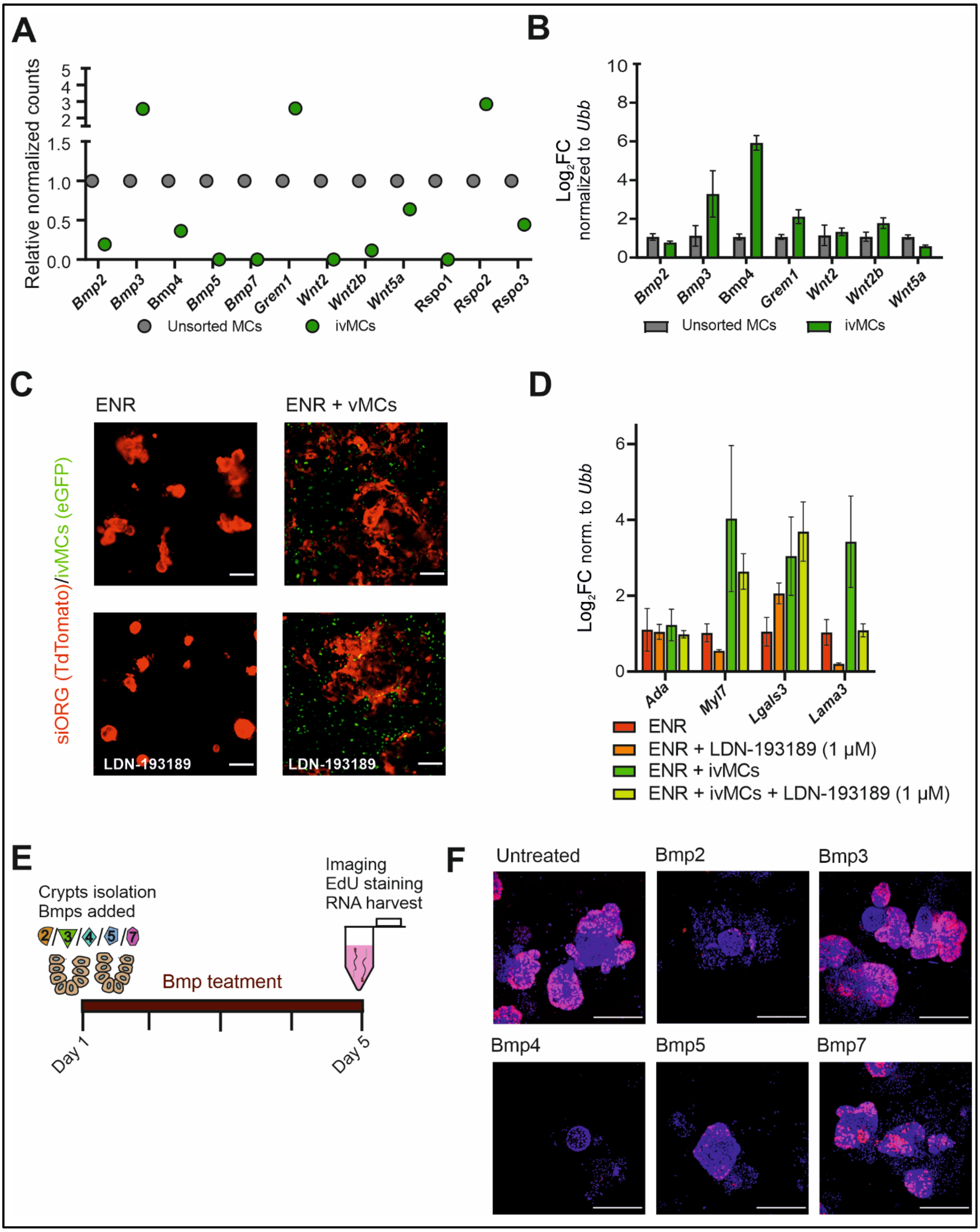
A
Relative normalized counts
Bmp2
Bmp3
Bmp4
Bmp5
Bmp7
Grem1
Wnt2
Wnt2b
Wnt5a
Rspo1
Rspo2
Rspo3
Unsorted MCs
ivMCs
B
Log$_2$FC normalized to Ubb
Bmp2
Bmp3
Bmp4
Grem1
Wnt2
Wnt2b
Wnt5a
Unsorted MCs
ivMCs
C
ENR
ENR + vMCs
siORG (TdTomato)/ivMCs (eGFP)
LDN-193189
LDN-193189
D
Log$_2$FC norm. to Ubb
Ada
Myl7
Lgals3
Lama3
ENR
ENR + LDN-193189 (1 μM)
ENR + ivMCs
ENR + ivMCs + LDN-193189 (1 μM)
E
Crypts isolation
Bmps added
Imaging
EdU staining
RNA harvest
Bmp teatment
Day 1
Day 5
F
Untreated
Bmp2
Bmp3
Bmp4
Bmp5
Bmp7

**Figure EV 1. Characterisation of ivMCs cells.**

A Immortalized cultured ivMCs still express Bmp ligands as determined by RNAseq. Dotplot indicates relative normalized counts, unsorted mesenchymal cells (MCs) were freshly isolated, short-time cultured mesenchymal cells from Pdgfra$^{H2BeGFP}$ animals, Time in culture: 9 weeks after the isolation from Pdgfra$^{H2BeGFP}$ animals, 7 weeks after the transduction with immortalisation retroviral particles, 3 weeks after sort/establishment of ivMCs.

B qRT-PCR of indicated Bmps and Wnts in vMCs after 6 weeks in the culture (total culture time since primary MSc isolation – 12 weeks). Expression levels normalized to *Ubiquitin B* (*Ubb*), n=3, unsorted freshly isolated MSCs parallel set as 1, error bars show sd.

C Co-cultivation of intestinal organoids (siORG, marked by tdTomato) with immortalized villus Mesenchymal cells (ivMCs, expressing nuclear eGFP) in the presence of Bmp type I receptor inhibitor LDN-193189 inhibitor (1 μM). ENR – organoids in cultivation medium (no vMCs), ENR + ivMCs organoids in cultivation medium with ivMCs. (Native fluorescent microscopy. Scale bar, 200 μm).

D Blocking Bmp type I receptor by LDN-193189 inhibitor (1 μM) partially revert villus-tip differentiation induced by vMCs as determined by qRT-PCR (Expression levels normalized to *Ubiquitin B* (*Ubb*), n=3, ENR parallel set as 1, error bars show sd).

E Scheme of 96 hours Bmp treatment of freshly isolated crypts.

F Proliferation of freshy isolated intestinal crypts cultivated with indicated Bmps for 96 hours determined by immunostaining for EdU incorporation (red), DAPI counterstains nuclei. (Immunocytochemistry, scale bar 200 μm).

*Figure EV 2*

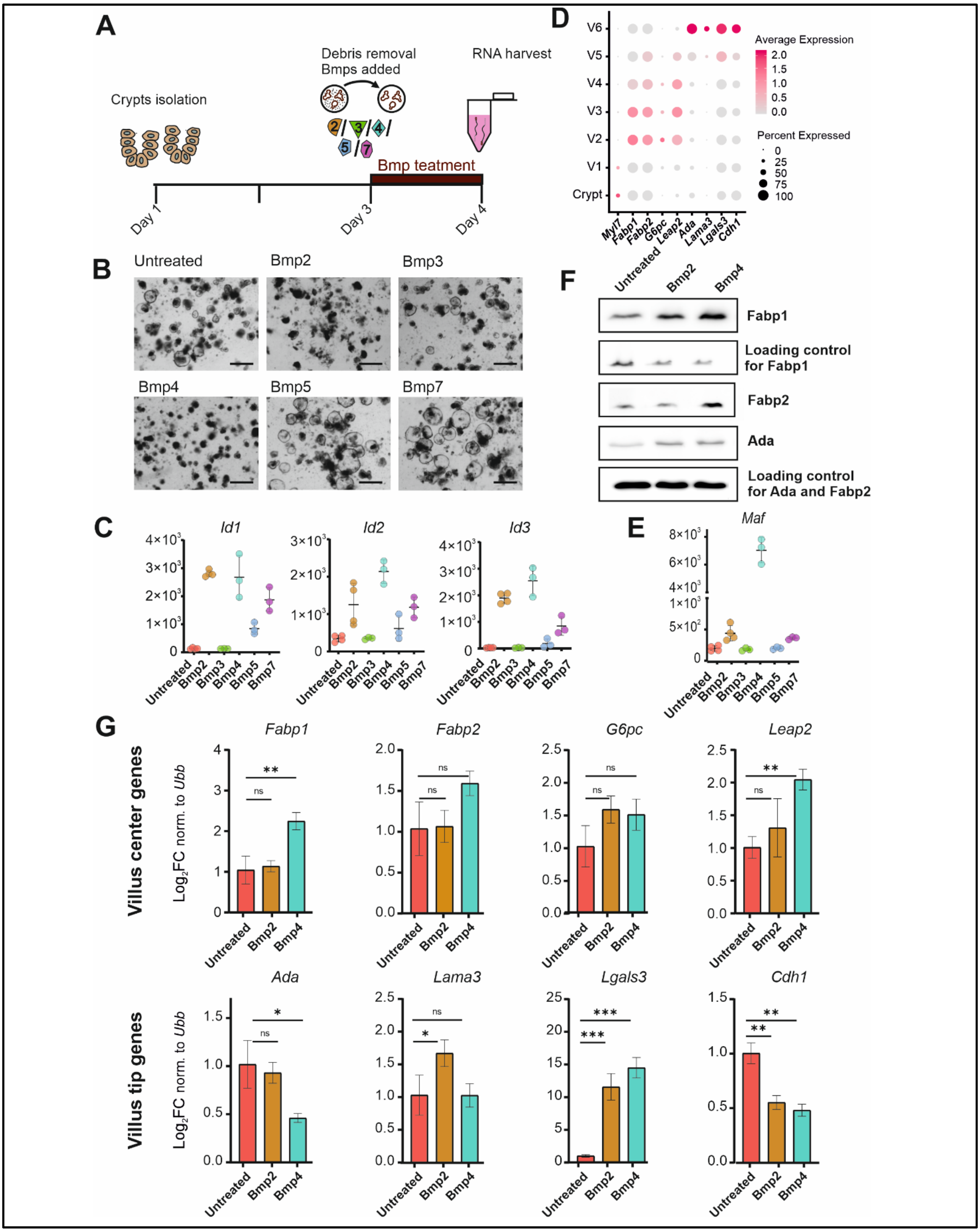

A
Crypts isolation
Debris removal
Bmps added
RNA harvest
Bmp teatment
Day 1
Day 3
Day 4
B
Untreated
Bmp2
Bmp3
Bmp4
Bmp5
Bmp7
C
Id1
Id2
Id3
D
Average Expression
Percent Expressed
E
Maf
F
Untreated
Bmp2
Bmp4
Fabp1
Loading control for Fabp1
Fabp2
Ada
Loading control for Ada and Fabp2
G
Villus center genes
Villus tip genes
Fabp1
Fabp2
G6pc
Leap2
Ada
Lama3
Lgals3
Cdh1

**Figure EV 2. Bmp2 induces villus tip programs in enterocytes, whereas Bmp4 activates lipid metabolism taking place in the center of the villus.**

A Scheme of 24 hours Bmp treatment of freshly isolated crypts.

B Brightfield image of freshly isolated crypts treated with 24 hours with indicated Bmp (Scale bar, 200 µm).

C Expression of generic Bmp target genes *Id1*, *Id2* and *Id3* upon 24 hrs Bmp treatment. (Graphs show normalized counts determined by RNAseq, n=3 for each treatment, error bars denote sd).

D The expression of indicated villus center genes and villus tip genes throughout the villus. (Dot plot, size of the dot represents the percentage of the cells expressing the transcript, color indicates the expression level. Used datasets: GSM2644349 and GSM2644350 reanalysed by Moor 2018: 10.5281/zenodo.3403670)

E The expression of *c-Maf*, the master regulator of lipid metabolism and uptake is initiated preferentially by Bmp4. (Graph shows normalized counts determined by RNAseq, n=3 for each treatment, time point: 24 hours error bars denote sd).

F Fabp2 protein connected to lipid uptake is expressed upon Bmp4 treatment. Immunoblot, of freshly isolated crypts treated for 48 hours by indicated Bmp2 (500ng/ml), Bmp4 (500ng/ml) or mixed Bmp2/4 (each 250 ng/ml), Fabp1 and Fabp2 are proteins involved in lipid uptake, Ada is an immunomodulator expressed at the villus tip. All proteins are expressed after Bmp-treatment, but Fabp2 is specifically regulated by Bmp4.

G qRT-PCR for villus center genes and villus tip genes in freshly isolated crypts cultivated 24 hours in Bmp2 (500ng/ml), Bmp4 (500 ng/ml) or mixture of Bmp2+Bmp4 (each 250 ng/ml). (Expression levels normalized to Ubiquitin B (*Ubb*), n=3, untreated parallel set as 1, error bars show sd, unpaired t-test, ns: $p > 0.05$, * $p \le 0.05$, ** $p \le 0.01$, *** $p \le 0.001$).

*Figure EV 3*

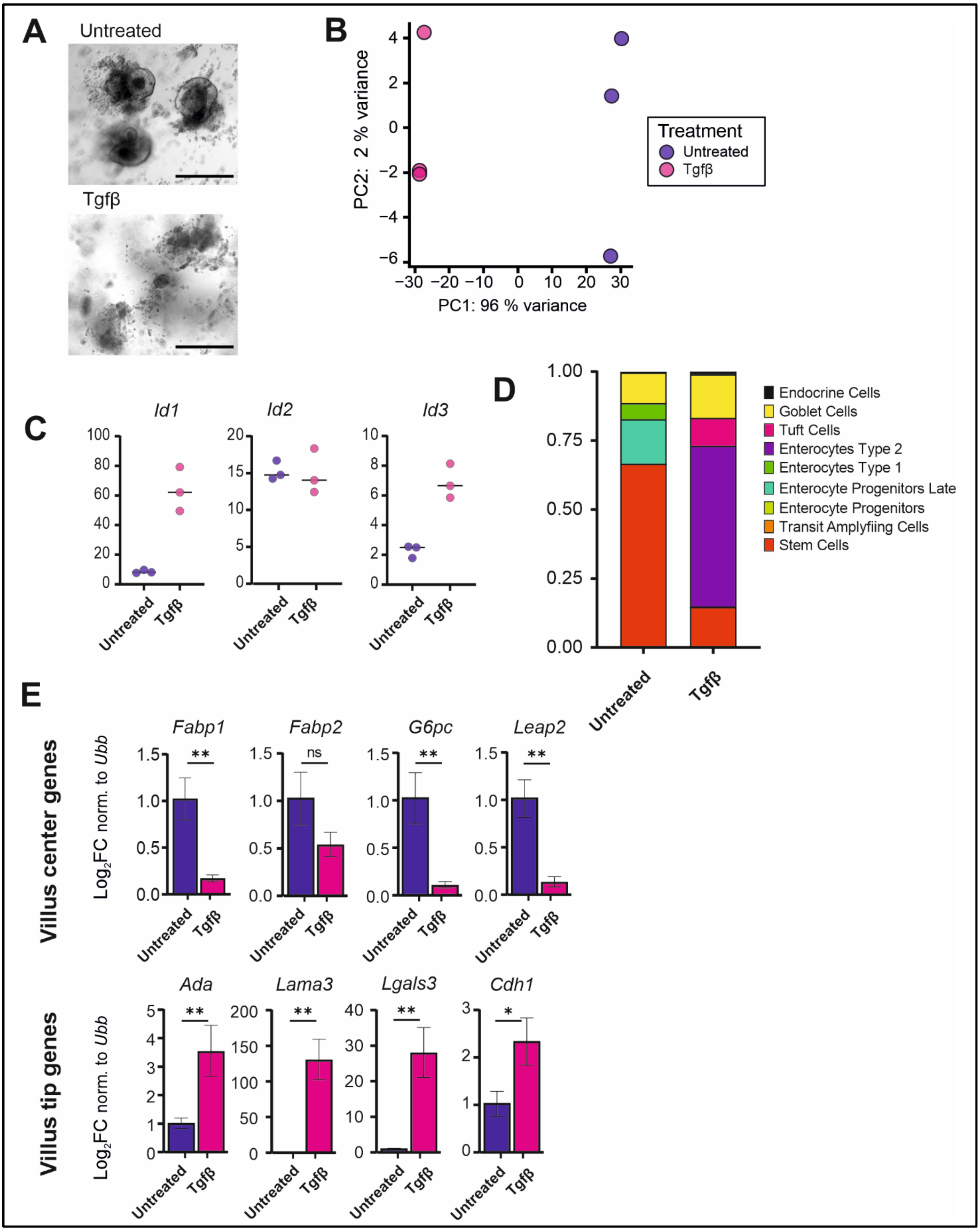

A
Untreated
Tgfβ
B
PC2: 2 % variance
PC1: 96 % variance
Treatment
Untreated
Tgfβ
C
Id1
Id2
Id3
Untreated
Tgfβ
D
Endocrine Cells
Goblet Cells
Tuft Cells
Enterocytes Type 2
Enterocytes Type 1
Enterocyte Progenitors Late
Enterocyte Progenitors
Transit Amplyfiing Cells
Stem Cells
E
Villus center genes
Fabp1
Fabp2
G6pc
Leap2
Villus tip genes
Ada
Lama3
Lgals3
Cdh1
Log2FC norm. to Ubb

**Figure EV 3. Tgfβ represses villus-center genes, but it activates villus tip-genes.**

A Brightfield image of freshly isolated crypts treated with 96 hours with Tgfβ. (Scale bar 200 μm)

B Principal component analysis (PCA) of the transcriptomes of intestinal crypts treated with Tgfβ for 24 hours. PC1 and PC2 explain 96% and 2% of the variance, respectively. (RNAseq samples, n=3 for each treatment and control).

C qRT-PCR for villus center genes and villus tip genes in freshly isolated crypts cultivated 24 hours with Tgfβ. Tgfβ villus center genes connected to lipid metabolism, while simultaneously activating expression of villus tip genes. (Expression levels normalized to Ubiquitin B (*Ubb*), n=3, untreated parallel set as 1, error bars show sd, unpaired t-test, ns= $p > 0.05$, * $p \leq 0.05$, ** $p \leq 0.01$, *** $p \leq 0.001$).

D Proportional changes of indicated epithelial cell types in the intestinal crypts treated with Tgfβ (500 ng/ml) for 24 hours. (Bar graph depicting deconvoluted RNAseq dataset compared by CellAnneal software to GSE92332 organoids scRNAseq dataset).

E qRT-PCR for villus center genes and vilus tip genes in freshly isolated crypts cultivated 24 hours in Tgfβ (500ng/ml). (Expression levels normalized to Ubiquitin B (*Ubb*), n=3, untreated parallel set as 1, error bars show sd, unpaired t-test, ns: $p > 0.05$, * $p \leq 0.05$, ** $p \leq 0.01$, *** $p \leq 0.001$).

*Figure EV 4*

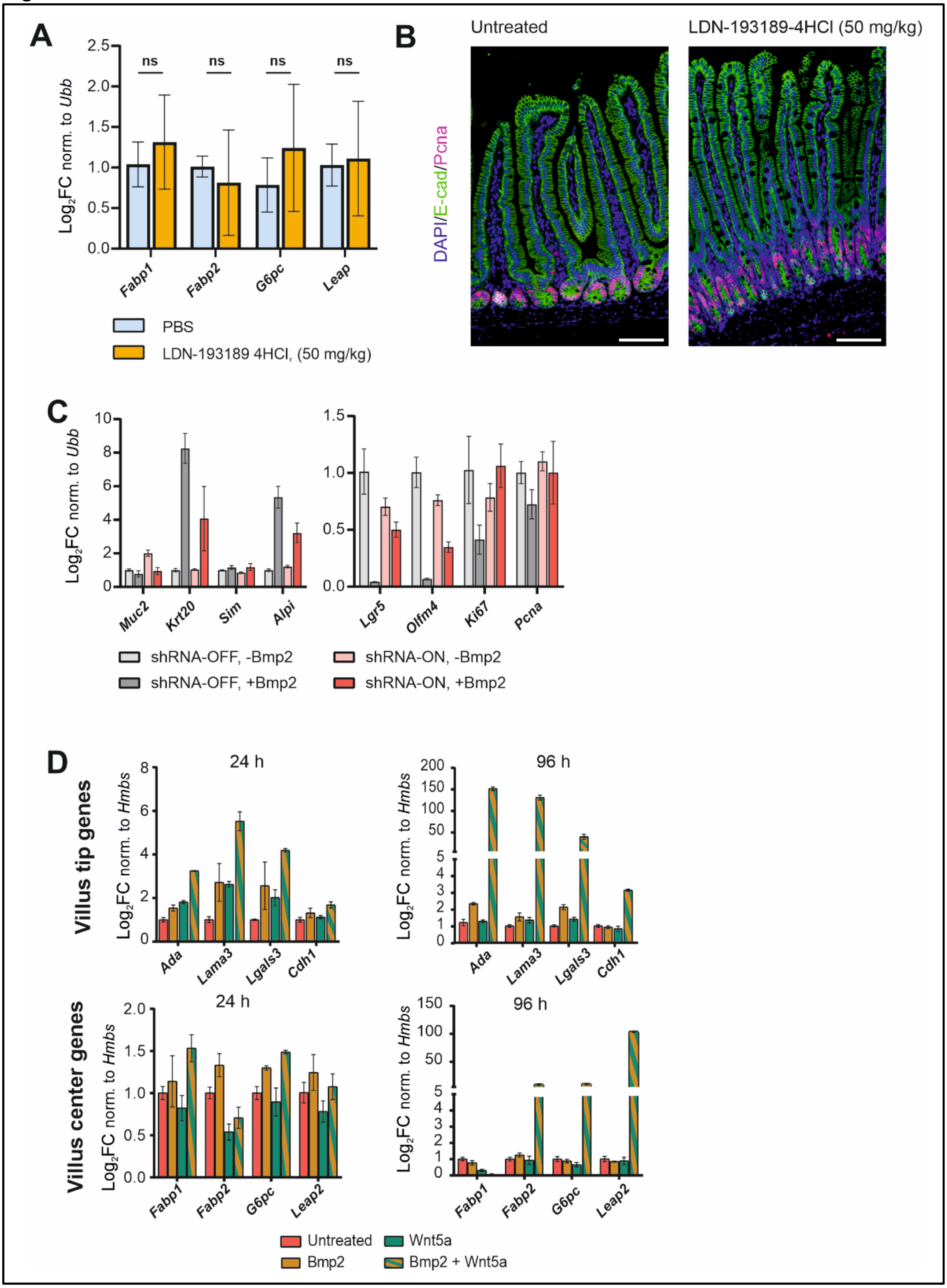

A
Log2FC norm. to Ubb
ns
Fabp1
Fabp2
G6pc
Leap
PBS
LDN-193189 4HCl, (50 mg/kg)
B
Untreated
LDN-193189-4HCl (50 mg/kg)
DAPI/E-cad/Pcna
C
Log2FC norm. to Ubb
Muc2
Krt20
Sim
Alpi
Lgr5
Olfm4
Ki67
Pcna
shRNA-OFF, -Bmp2
shRNA-OFF, +Bmp2
shRNA-ON, -Bmp2
shRNA-ON, +Bmp2
D
Villus tip genes
24 h
96 h
Log2FC norm. to Hmbs
Ada
Lama3
Lgals3
Cdh1
Villus center genes
Fabp1
Fabp2
G6pc
Leap2
Untreated
Wnt5a
Bmp2
Bmp2 + Wnt5a

**Figure EV 4. Despite Bmp2 induces villus-center programs in canonical Smad-dependent manner by its own, it acts synergistically with Wnt5a.**

A Four days treatment with LDN-193189 inhibitor does not significantly affect the expression of the villus-center genes. (qRT-PCR, expression levels normalized to *Ubiquitin B* (*Ubb*), n=3, error bars show sd, untreated parallel set as 1).

B The normal epithelial morphology and proliferation after the 4 days treatment with LDN-193189 inhibitor. Immunofluorescence, E-cadherin (green) marks epithelial cells, Pcna (red) stains proliferating cells, DAPI (blue) counterstains nuclei. Scale bar 200 μm

C Smad4 knock-down (shRNA-ON) alleviates Bmp2-induced differentiation marked by relative expression of differentiation markers (*Muc2, Krt20, Sim, Alpi*) and restores the expression of stem cell and proliferation genes (*Lgr5, Olfm4, Ki67, Pcna*). (qRT-PCR, expression levels normalized to *Ubiquitin B* (*Ubb*), n=3, error bars show sd, untreated parallel set as 1, treatment as in Figure 4.C,D).

D qRT-PCR for villus tip genes and villus center genes in freshly isolated crypts cultivated 24 or 96 hours with Bmp2 (500 ng/ml), Wnt5a (500 ng/ml) or both Bmp2 + Wnt5a (each 500 ng). Whereas Bmp2 induces villus tip genes by its own, it can act synergistically with Wnt5a. For the short-term cultivation, the effect is specific for villus-tip genes (notice the different scale in left graphs). (Expression levels normalized to *Hmbs*, n=3, untreated parallel set as 1, error bars show sd).
